# Thermohydraulic Effect of Zebra Stripes - Numerical Analysis

**DOI:** 10.1101/2023.05.02.539143

**Authors:** Vojtěch Smolík

## Abstract

The function of zebra stripes has many possible explanations. Some researchers suggested zebra stripes are a form of adaptation to high temperatures, allowing to maintain a lower animal body temperature in hot environment. Sunlight radiation creates a temperature difference between cooler white and hotter black stripes. Hypothesis state there are currents of air formed caused by this temperature difference. This hypothesis is examined using the CFD numerical simulation of zebra surface in ANSYS Fluent to uncover the physical phenomenon.

## 1. Introduction

Zebra stripes are a natural phenomenon that is not fully understood. In the article *”How the zebra got its stripes: a problem with too many solutions”* [1] researchers explain many possible explanations as social cohesion, thermoregulation, predation evasion and avoidance of biting flies. This paper states the pattern of zebra stripes is in strong correlation with environment temperature. The Figure 1 shows a distribution of different zebra species in Africa. Zebras in hot areas near the equator tend to have more significant stripe patterns with thicker and more contrast stripes. However, this finding is a weak evidence for any conclusion, since the number of environmental variables is high. The thermoregulation hypothesis says: sunlight radiation creates an uneven distribution of temperatures on the surface of zebra body. Black stripes are heated up to a higher temperatures thanks to the higher emissivity compared to white stripes. The temperature difference creates a natural convection currents flowing near the body surface: increasing the evaporation rate of sweat and maintaining lower body temperature.

**Figure 1:**
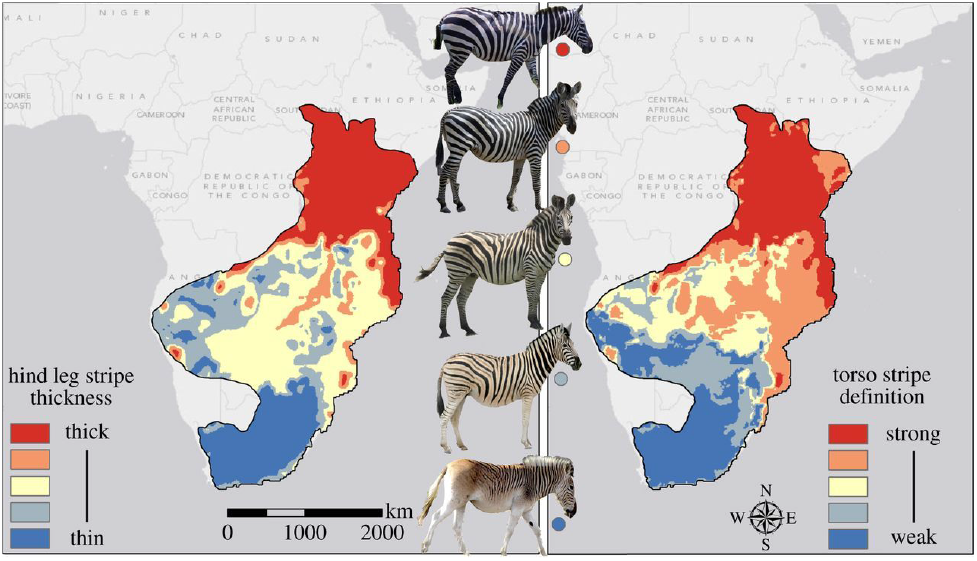
Predicted levels of hind leg stripe thickness (left) and torso stripe definition (right), from a random forest model based on 16 populations. [1]

This paper is focused only on the physical phenomenon of the thermoregulation hypothesis, other biological factors are neglected. Horváth article [2] presents the idea of air flow near the zebra surface, shown in the Figure 2(a). This air flow mechanism can be also influenced by the effect observed by Alison and Stephen Cobb [3]: zebras tend to errect black hairs over the white ones. The difference of height between white and black stripes (piloerection) is shown in the Figure 2(b).

**Figure 2:**
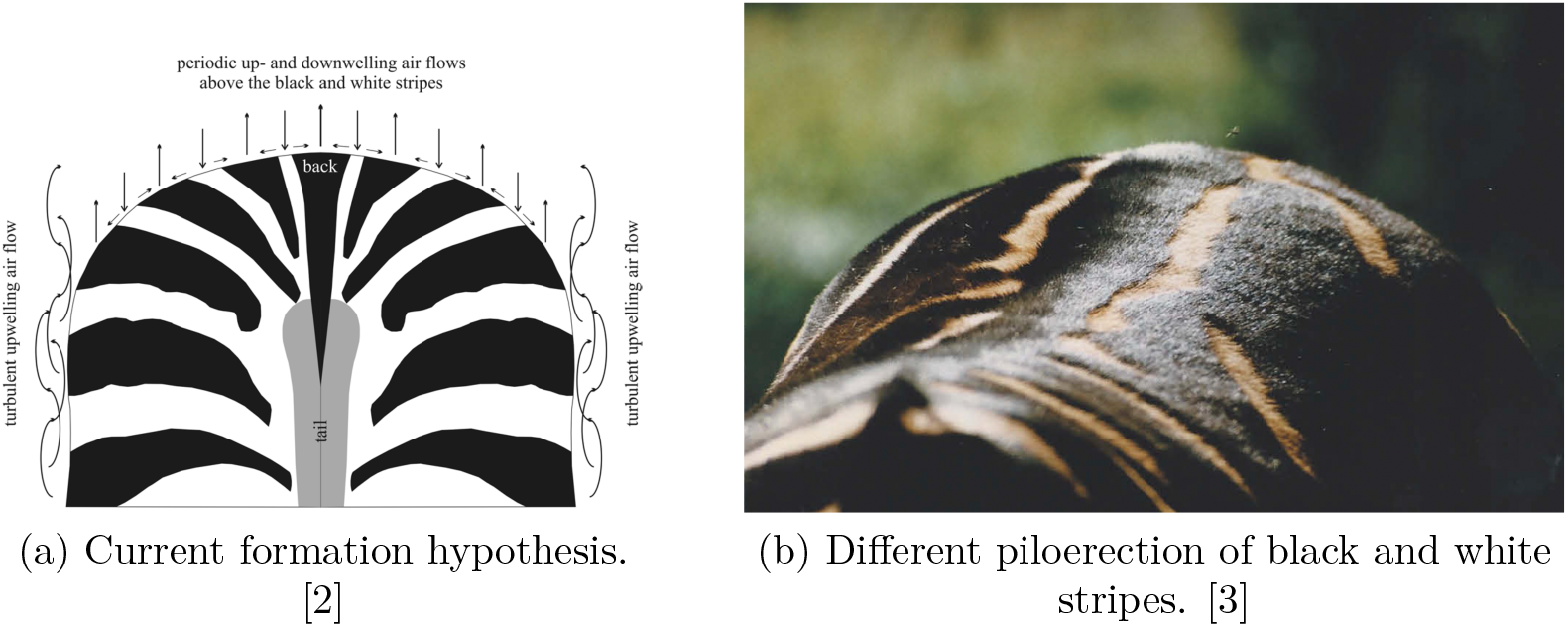
The possible mechanisms of the thermoregulation hypothesis.

## 2. Methods

Boundary conditions of the numerical model are average temperatures of black and white stripes and air temperature measured by Alison and Stephen Cobb, shown in the Figure 3. Temperatures taken from Figure 3 at 12:00 are 55 °*C*, 40 °*C* and 26 °*C*. Authors measured the temperatures with a hand-held digital infra-red thermometer.

**Figure 3:**
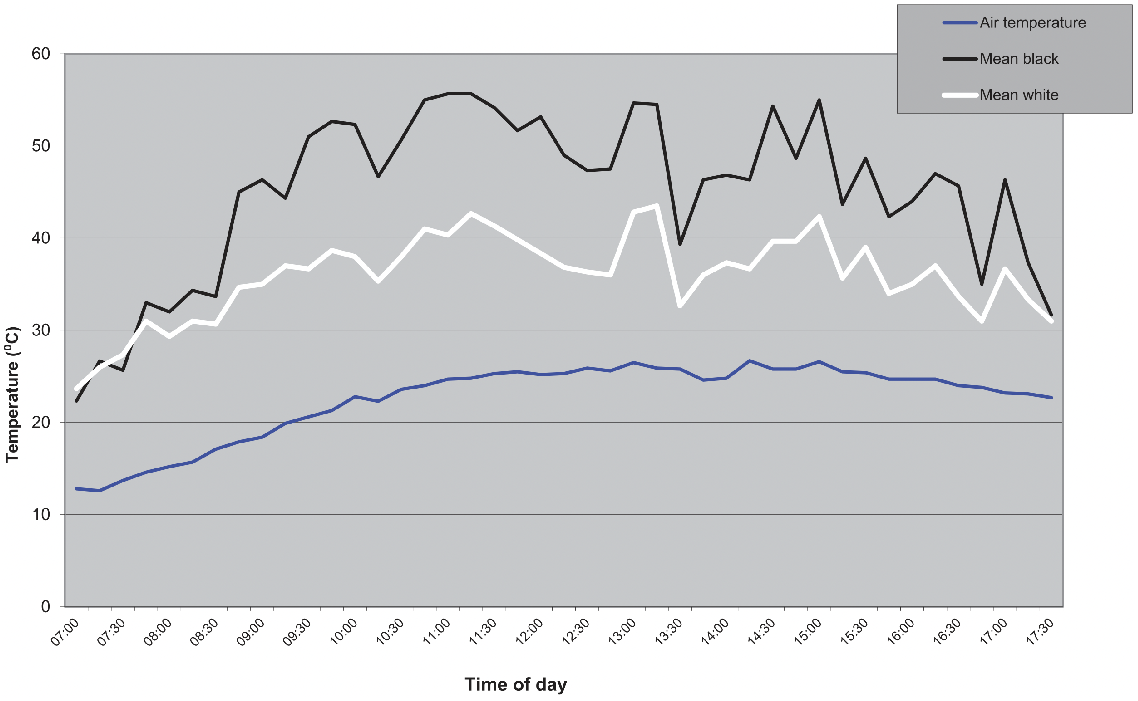
Zebra mare stripe temperatures. [3]

Two parameters were studied in the numerical analysis: the temperature difference between stripes and the black stripe height effect. The total of 6 models were simulated as a combination of selected parameters, shown in the Table 1. The Figure 4 shows simplified zebra pattern geometry used for numerical model. Only a selected part was used for simulation to reduce the complexity of model. The temperatures of zebra surface are set as boundary condition to a constant value. The goal of numerical analysis is to examine the air flow and air temperature near the animal body surface.

**Table 1:**
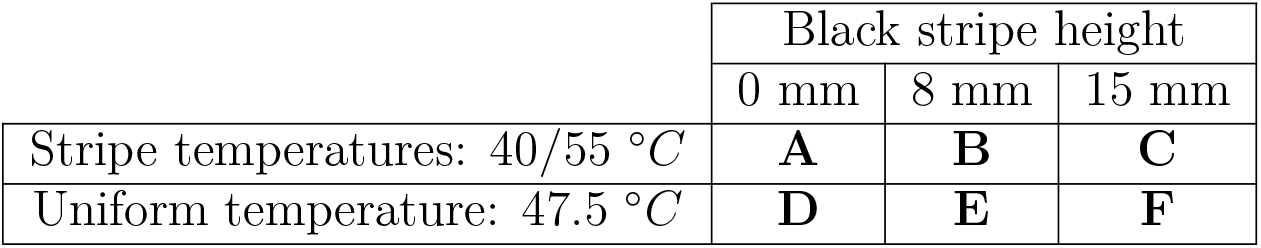
Numerical Analysis Models

|  | Black stripe height |  |  |
| --- | --- | --- | --- |
|  | 0 mm | 8 mm | 15 mm |
| Stripe temperatures: 40/55 °C | **A** | **B** | **C** |
| Uniform temperature: 47.5 °C | **D** | **E** | **F** |

**Figure 4:**
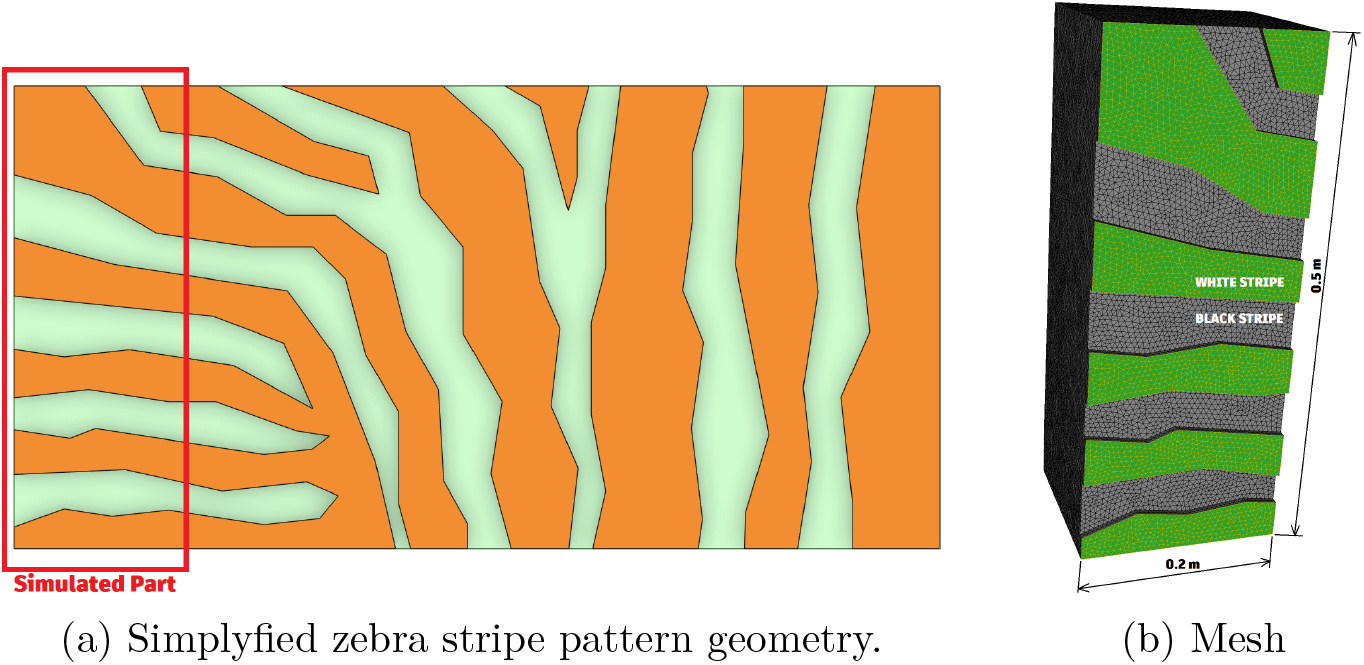
Zebra geometry.

Boundary walls surrounding the simulated volume are set as pressure-outlet to allow free movement of air. Air velocity in the model was set to zero to not disrupt the current creation. The effect of outside air flow around the zebra body was examined separatelly. Gravity and temperature dependent density of air were considered to create the natural convection phenomenon. In the second part of numerical simulation, the air flow in the saddle area of zebra was examined by turning the gravity acceleration perpendicular to the body surface. The example of mesh used for numerical simulation is shown in the Figure 4(b).

## 3. Results

A total of 3 sets of numerical simulations were performed. The first one studied the effect of zebra stripe temperature difference and black stripe height. Zebra stripe pattern with a temperature difference of 15°*C* (55-40 °*C*) is compared with uniform temperature surface (47.5 °*C*). Black stripe height was set to 0, 8 and 15 mm. The largers value is exaggerated to show the mechanisms of air flow. The second set of simulations examined the influence of outside air flow velocity. The velocity of air (flowing in parallel to zebra stripes) was set to 0, 0.5 and 1 m/s. The last set of simulations focused on the air behaviour in the saddle area of zebra. This was accomplished by changing the direction of the gravitational acceleration.

Table 2 shows the first set of numerical analysis results. Air flow streamlines are colored according to the local air temperature. First row corresponds to the zebra pattern surface with a temperature difference between stripes. The second row represents samples with uniform temperature surface. Sample A shows no significant air flow patterns near the body surface compared to sample D. Temperature difference between stripes affected the local temperature of air, however it caused only minor changes in the air flow.

**Table 2:**
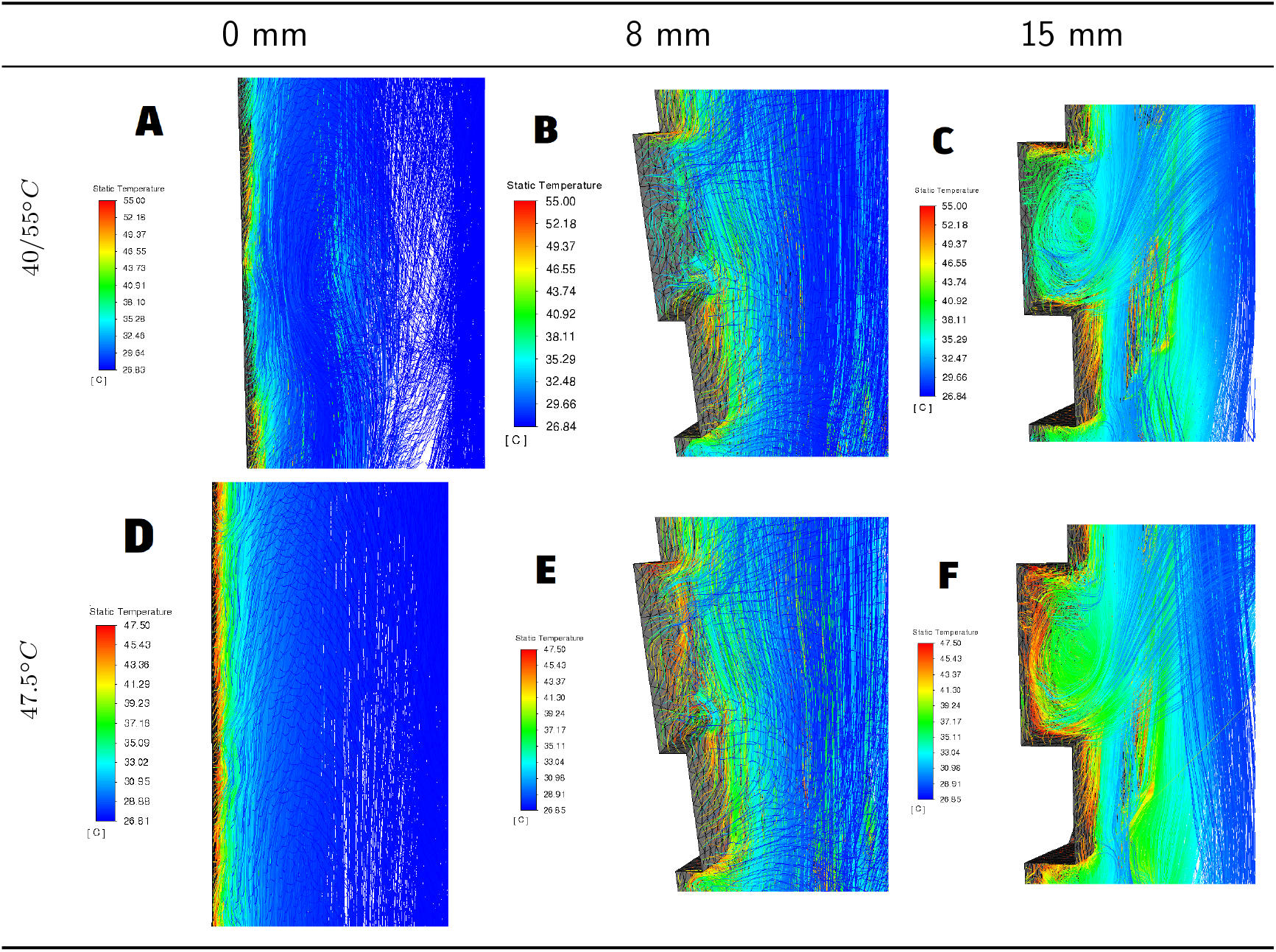
The results of numerical analysis.

Figures B and E show the disturbance of air flow caused by the black stripe thickness. There are no significant differences between samples B and E, thus the effect of stripe thickness on the air flow is substantially stronger than the effect of temperature difference between stripes. Figures C and F show the exaggerated sample with 15mm black stripe thickness. This sample shows how the air flow will theoretically evolve in such case. Table 3 shows the second set of numerical results. The effect of outside air flow velocity on the natural convection near the animal body is studied. The sample C with zebra pattern and largest black stripe thickness performed the most significant natural convection air flow, thus it was selected for this simulation. The results in Table 3 show the natural convection air flow patters are distubed even by the lowest 0.5 m/s outside air flow velocity.

**Table 3:**
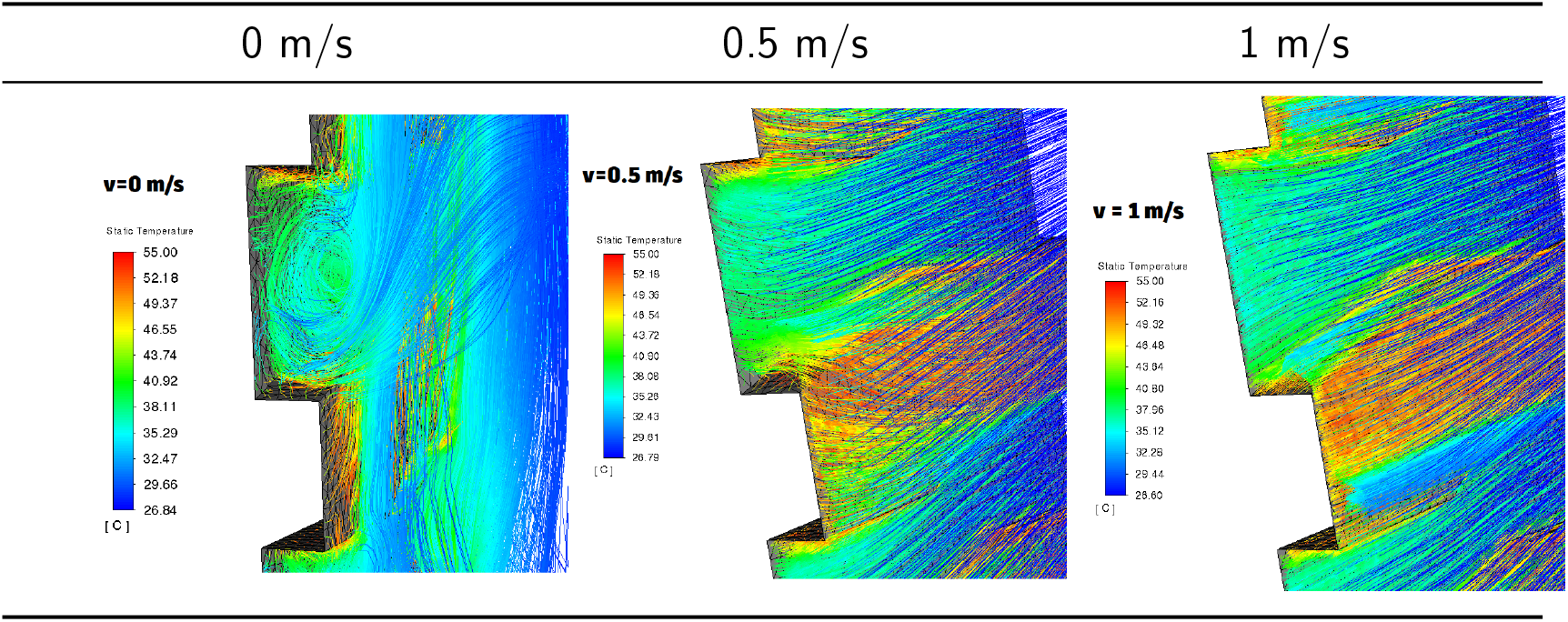
The effect of external air flow velocity applied on sample C.

The last set of results shows the natural convection air movement in the saddle area of zebra. Figure 5 shows the air flow with corresponding air temperatures over the zebra striped flat surface. Regions above the black stripes with higher temperature reach higher air temperatures. The natural convection air flow differs from the sample with uniform surface temperature, shown in the Figure 6. However, the zebra stripe pattern caused no significant air flow pattern beneficial for heat transfer.

**Figure 5:**
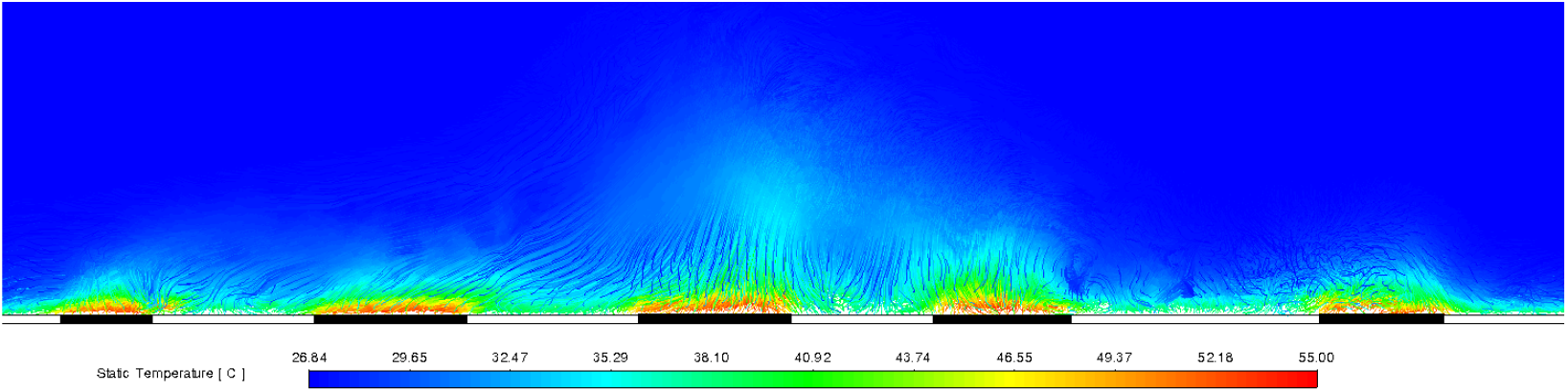
The saddle area air flow - zebra stripe pattern (40*/*55°*C*).

**Figure 6:**
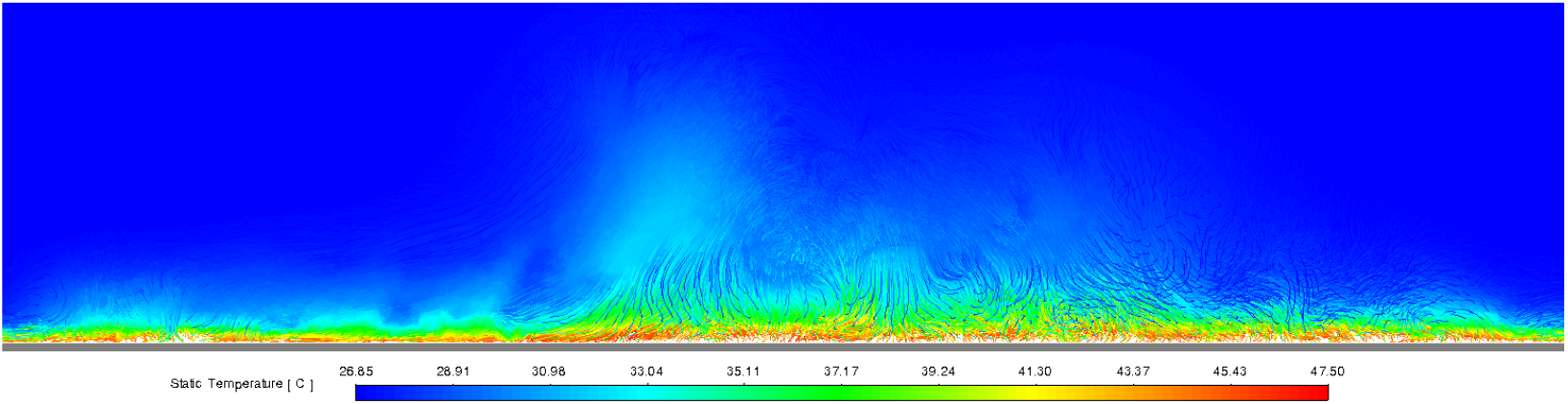
The saddle area air flow - gray surface with uniform temperature (47.5°*C*).

All numerical analysis were performed on tetrahedral type mesh. The dimensions of simulated volume are 0.2×0.2×0.5 m, shown in the Figure 4(b). The temperature dependent density of air was set as a piecewise linear function. Realizable k-epsilon viscous model was used for the simulation.

## 4. Discussion

Results of numerical analysis showed no direct proof of the thermoregulation effect hypothesis based on the temperature difference between stripes. However, the piloerection of black stripes resulted in the formation of convective current patterns near the animal body surface. Those currents are beneficial for the animal thermoregulation, since they are *”a reliable medium for air and water vapour exchange during sunlight, thus facilitating the removal of air saturated with sweat*.*[3]”* Alison and Stephen Cobb observed the piloerection effect (increasing the black stripes) is changing in the course of the day. Zebras tend to erect black stripes in the morning hours, but also in the hottest time of the day. The future research focused on the correlation between the piloerection and the environment temperature is strongly suggested. Alison and Stephen Cobb: *”As far as we are aware, this differential piloerection has not been commented on previously*.*” [3]*

## 5. Conclusion

This article focused on the zebra stripe thermoregulation effect hypothesis. A total of 11 numerical simulation results were presented to examine the impact of zebra stripes on the natural convection around the animal body surface. Physical models studied the effect of temperature difference between black and white stripes and the effect of black stripe thickness. It was found that the temperature difference between stripes does not have a significant role in natural convection streams formation. However, the thickness of black stripes exceeding the white ones can result in formation of various air currents with potential to enhance the animal surface thermoregulation function. Zebras errect black stripes differently in the course of the day. This phenomenon might help to prove the thermoregulation function of variable stripe thickness, since the black stripes are observed to be erected in the hottest hours of the day. The study of correlation between the piloerection effect and zebra body thermoregulation is suggested for a further research.

